# The Omega-3-fatty acid docosahexaenoic acid modulates intracellular *myo*-inositol and its biosynthetic genes

**DOI:** 10.1101/2021.08.25.457630

**Authors:** Marlene Murray, Taejun Ok, Haley Kang, Jee Yeon Lee, Bomi Kim, Hyukje Sung, Talisa Tait, Daniel Colon Hidalgo, Yudy Guzman

## Abstract

Bipolar disorder is a debilitating mood disorder characterized by recurring episodes of mania and depression. It affects 2.6% of adults and has a lifetime prevalence among adults of 3.9%. Current mood stabilizers are not always effective and/or are not well tolerated by many patients; thus, there is a need to develop or identify more effective and less harmful treatments. Omega-3-fatty acids have been shown to be effective in the treatment of bipolar disorder; however, their mechanism of action is unknown. *Myo*-inositol depletion has been hypothesized as the mechanism by which mood stabilizers exert their therapeutic effect. Using an enzymatic assay, we determined intracellular *myo*-inositol levels increased more than 2-fold when cells were grown in the presence of the omega-3 fatty acid docosahexaenoic acid (DHA). RT-qPCR was used to characterize the effects of DHA on genes in the *myo*-inositol biosynthetic pathway. We show DHA increases relative expression and has a concentration-dependent impact on *INO1* and *INM1*, which encode *myo*-inositol-1-phosphate synthase and *myo*-inositol monophosphate 1-phosphatase, respectively. We therefore conclude that the omega-3-fatty acid DHA exerts its therapeutic effect on bipolar disorder by increasing intracellular *myo*-inositol, which may be accomplished by upregulating its biosynthetic genes.

## Introduction

Omega-3-fatty acids are long-chain polyunsaturated fatty acids (PUFA) derived from plants and marine sources. They are essential and must be derived from dietary sources. The three main omega-3-fatty acids are alpha-linoleic acid (ALA), eicosapentaenoic acid (EPA) and docosahexaenoic acid (DHA). While EPA and DHA are found in fish and other seafood, ALA is mainly plant-derived and is a precursor to EPA and DHA. Conversion of ALA to EPA and DHA is not efficient in humans so adequate levels can only be achieved via food consumption (Burdge, 2006).

Omega-3-fatty acids are integral to cell membranes and are known to affect membrane receptors. They provide the building blocks for hormone synthesis leading to downstream effects on human physiology. Not surprisingly, it has been shown consumption of foods containing these fatty acids correlates with a lower prevalence of bipolar disorder (Noaghiul and Hibbeln, 2003). Bipolar disorder is one of the most debilitating and severe mood disorders. It is characterized by recurring episodes of mania and depression, with the possibility of each episode lasting up to several months. According to the WHO’s most recent study on the global burden of disease, bipolar disorder ranks within the top 20 causes of medical conditions worldwide and 6^th^ among mental disorders. Omega-3 fatty acids have been used as adjuvants to bipolar disorder treatments such as lithium and valproate (VPA), resulting in reduced symptoms, specifically depression and reduced relapse rates (Krawczyk and Rybakowski, 2012; Logan, 2003; Nemets et al, 2002; Frangou et al., 2006; Osher, 2005; Stoll et al., 1999; Ross et al., 2007). They are safe and are not associated with the harmful effects seen with standard treatments. While the molecular mechanism underlying the therapeutic benefit of omega-3 fatty acids is unknown, there is evidence that like lithium and VPA; they affect the phosphatidylinositol signaling pathway. Specifically, they have been shown to inhibit protein kinase C (PKC), phospholipase C (PLC) and inositol triphosphate (IP_3_), all of which are known to be affected by VPA and lithium (Mirnikjoo et al., 2001; Seung Kim et al., 2001; Sarkar et al., 2005; Sperling et al., 1993). Not surprisingly, this pathway is also implicated in the pathophysiology of the disease. Jope et al., 1996 reported decreased activity of the phosphoinositide signal transduction system in the occipital cortex of bipolar victims obtained postmortem while Brown et al., 1993 and Soares et al., 2001 reported elevated phosphatidylinositol-4,5-bisphosphate (PIP_2_) levels in platelet membrane of both bipolar mania and drug-free bipolar patients in the depressed state. Additionally, altered levels of protein kinase A (PKA), and Rap1 (PKA substrate) have been observed in several regions of the brains of bipolar disorder patients, and increased phosphorylation of Rap1 has also been detected (Perez et al., 1999). The activity of platelet PKC has been found to be elevated during manic episodes (Friedman et al., 1993).

The depletion of *myo*-inositol has been proposed as the therapeutic mechanism that alleviates the symptoms of bipolar disorder. This hypothesis though controversial, is supported by the fact that both lithium and VPA decrease *myo*-inositol and affect other components of the phosphatidylinositol signaling pathway. These effects are reversible upon the addition of *myo*-inositol. (O’Donnell et al., 2000; Shaltiel et al., 2004). *Myo*inositol is a simple six-carbon sugar that serves as the precursor to the inositol containing phospholipid-phosphatidylinositol and as a backbone to several important intracellular signaling molecules. Therefore, drugs that interfere with the formation of these compounds, as is the case of both lithium and VPA, may have a significant impact on a wide range of cellular functions, some of which may be associated with bipolar disorder. In fact, postmortem brain analysis of suicide victims and patients with bipolar disorder reveal low levels of *myo*-inositol in the frontal cortex (Shimon et al., 1997). In addition, a comparison of the metabolites in the frontal brain of depressive patients and normal individuals indicate lower *myo*-inositol levels in depressive patients (Gruber et al., 2003) and increased levels in the left dorsolateral prefrontal cortex during manic episodes of bipolar patients (Frey et al., 2005). Previous studies have employed *myo*-inositol depletion in yeast as a screening tool to identify potential new anti-bipolar treatments (Ding et al., 2009). In the current study, we used the yeast *Saccharomyces cerevisiae* as a model system to determine the effects of the omega-3-fatty acid DHA on intracellular *myo*-inositol levels and expression of genes in the *myo*-inositol biosynthesis pathway. The genes are *INO1* and *INM1. INO1* encodes *myo*-inositol-1-phosphate synthase, which catalyzes the conversion of glucose-6-phosphate to *myo*-inositol-1-phosphate. This is the rate-limiting step in the de novo synthesis of *myo*-inositol. The *INM1* gene encodes *myo*-inositol monophosphate 1-phosphatase, which catalyzes the conversion of *myo*-inositol-1-phosphate to *myo*-inositol. Here we report that DHA increases intracellular *myo*-inositol and expression of both *INO1* and *INM1*.

## Materials and Methods

### Strain

The *Saccharomyces cerevisiae* strain used in this study was SMY7 (derivative of D273-10B/A1, *ade5*, *ura3*, *MAT a*).

### Media and Growth Conditions

Cells were cultured in complete medium [0.069% vitamin-free yeast base, 2% glucose, 0.201% ammonium sulfate, 20mg/l adenine, 20mg/l arginine, 10mg/l histidine, 60mg/l leucine, 20mg/l lysine 20mg/l methionine, 300mg/l threonine, 20mg/l tryptophan, 40mg/l uracil, and vitamins (described in Culbertson and Henry, 1975)], with the indicated concentrations of DHA and VPA as a positive control. The concentrations of DHA and VPA used were determined from previous experiments (data not shown) as the highest concentrations that permitted growth. Cultures were inoculated to an A_550_ of 0.1 and grown for 24 hours at 30°C in a shaking water bath at 150 rpm.

### Cell extracts

Cultures were centrifuged, and pellets were washed twice before being resuspended in deionized water to a 1mg/ml concentration. Glass beads were added to ~50% of volume, and the mixture vortexed for 10 minutes at 2-minute intervals, alternating with 2-minute incubations on ice. Extracts were clarified by centrifugation for 5 min at 2,000 rpm. Supernatants were collected and stored at −80°C. The protein concentration of the extracts was determined by the Bradford (1976) method using bovine serum albumin as the standard.

### Measurement of intracellular myo-inositol

*Myo*-inositol in cell extracts was measured by the enzyme-spectrophotometric assay of Ashizawa *et al*. (2000). Cell extracts were deproteinized with 16% (w/v) perchloric acid and centrifuged at 5,000 rpm for 10 minutes. The resultant supernatant was neutralized with 2M K2CO3 before being dephosphorylated with hexokinase reagent (200 mM Tris-HCl buffer, 400 mM adenosine triphosphate disodium (pH adjusted to 8.6 with 10M NaOH) and 115U/ml hexokinase) then incubated at 37°C for 90 minutes. The reaction was stopped by heating for 3 minutes in a boiling water bath then 4.5 M HCl was added to remove endogenous NADH and NADPH. After 10 minutes at 25°C, 3M K_2_CO_3_ was added for neutralization. The neutralized extract was then mixed with *myo*-inositol reagent (210mM triethanolamine hydrochloride-32mM K_2_HPO_4_-KOH buffer (pH 8.6), 1.2% (v/v) Triton X-100, 10mM β-NAD, 1.0 U/ml diaphorase, 0.1% (w/v) bovine serum albumin, 60μg/ml INT). The absorbance of the solution was measured at 492 nm with a microplate reader and the reaction initiated by addition of 2.1U/ml *myo*-inositol dehydrogenase dissolved in 20 mM potassium phosphate buffer (pH 7.0) to each well. After 30 minutes at room temperature, absorbance at 492 nm was again measured. *Myo*-inositol content was calculated from ΔA as an increase in absorbance during the reaction. Concentration of *myo*inositol in the cell extracts was determined using a standard curve which was linear in concentrations ranging from 0 mM to 10 mM.

### RNA extraction and cDNA synthesis

Total RNA was extracted from cells using the Aurum^™^ Total RNA Mini Kit (Bio-Rad). The spin protocol was employed according to the manufacturer’s instructions. RNA concentration was measured using a spectrophotometer in 100μl volume. cDNA synthesis was performed using iScript Reverse Transcription Supermix for RTqPCR (Bio-Rad) according to the manufacturer’s instructions with reverse transcription initiated by oligo (dT)-random primers supplied with the kit.

### RT-qPCR

The relative expression of *INO1* and *INM1* were compared to that of the reference gene *TFC1* (house-keeping gene). RT-qPCR reactions of 20μl volume contained the Sso Advanced^™^ Universal SYBR^®^ Green Master-Mix and primers for SYBR Green assays, both obtained from Bio-Rad. Reactions were performed on the Bio-Rad^®^ CFX96^™^ system under the following conditions: 30 s at 95°C for polymerase activation and 40 cycles of 5 s at 95°C and 30 s at 62°C. Melting curves were obtained between 65 to 95°C with a ramp rate of 0.5°C/s. The data presented were normalized to *TFC* 1 mRNA. To determine the relative mRNA expression, we used the delta-delta CT (2^-ΔΔCT^) method (Livak and Schmittgen, 2001).

### Statistical analysis

The significance of the differences between the means of intracellular inositol concentration and relative mRNA expression was determined by One-way ANOVA. A p-value < 0.05 was considered significant.

## Results

### Effect of DHA on intracellular myo-inositol

Previous studies showed that VPA decreased intracellular *myo*-inositol (Vaden *et al.*, 2001). Figure 1 shows 0.2 mM DHA decreased *myo*-inositol by 3.1-fold but resulted in an approximately 2.3-fold increase when cells were grown in 0.4 and 0.6mM DHA. The differences were statistically significant (*F* _4,23_= 4.33, *p* < .05).

**Figure 1.**
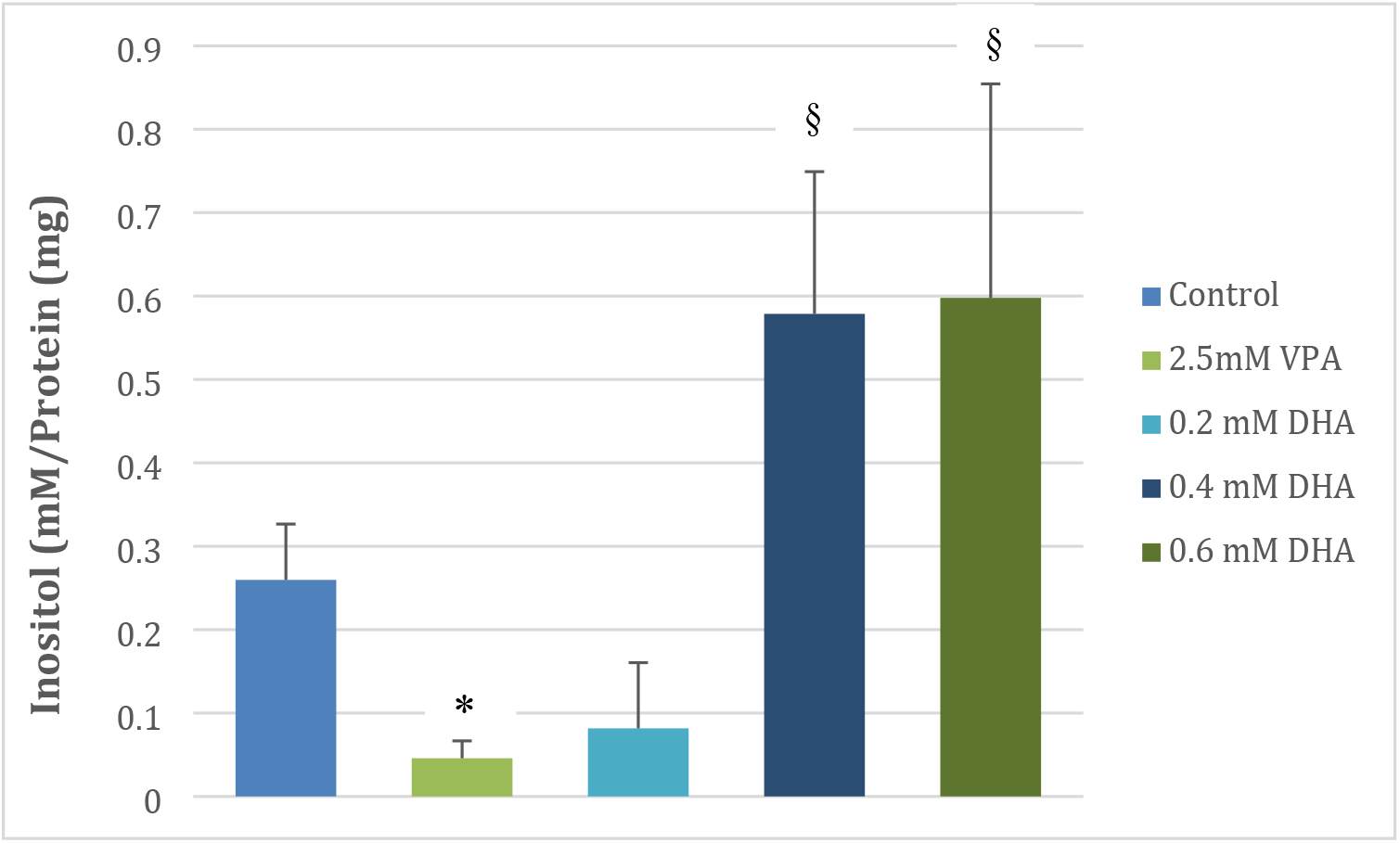
Mean ± SE of intracellular *myo*-inositol concentration in extracts of cells grown in the absence (control, n= 8) or presence of the indicated concentrations of docosahexaenoic acid (DHA), n= 2-7, and valproate (VPA), n= 8. * Significantly different from control at p-value < 0.05. § Significantly different from VPA at p-value < 0.05

### Effect of DHA on INO1 gene expression

Previous studies showed VPA increased *INO1* gene expression (Vaden *et al.*, 2001). Figure 2 shows the relative expression of *INO1* mRNA by RT-qPCR. The relative expression of *INO1* displayed an inverse relationship with DHA concentration. Expression was 1.4, 1.2 and 0.8 in the presence of 0.2mM, 0.4mM and 0.6mM DHA respectively. The differences observed were not significant (*F* _3,21_=1.23, ns).

**Figure 2.**
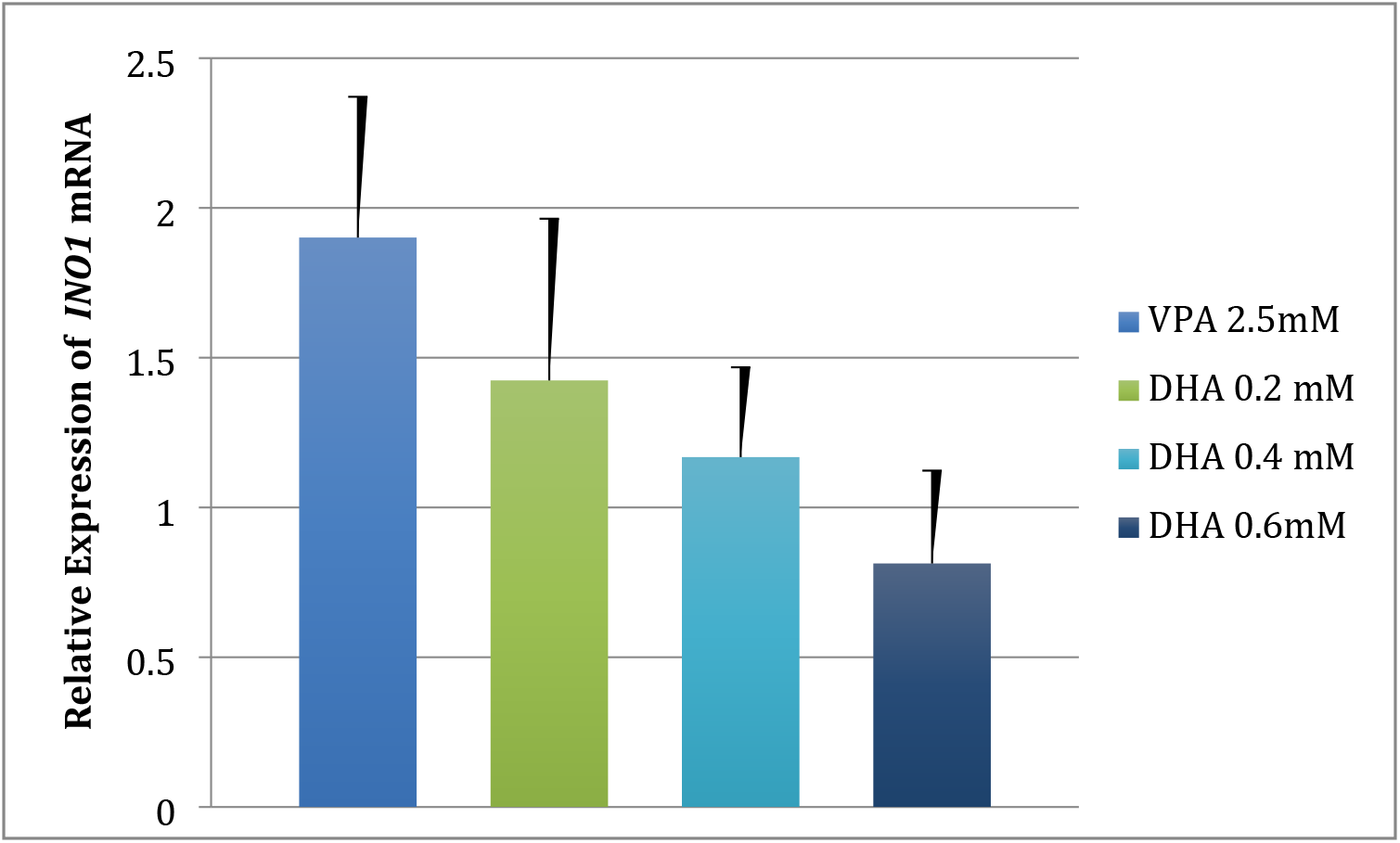
Relative *INO1* gene expression of cells grown in DHA (n =6) and positive control (VPA, n =7). The delta-delta CT method was used to determine fold change. *p* > 0.05 by ANOVA.

### Effect of DHA on INM1 gene expression

Figure 3 shows the relative expression of *INM1* mRNA by RT-qPCR. DHA resulted in overexpression of *INM1* with expression increasing 1.5-fold at the highest DHA concentration (0.6mM). The differences were not statistically significant, (*F* _3,16_ = 4.29, *p* < .05)

**Figure 3.**
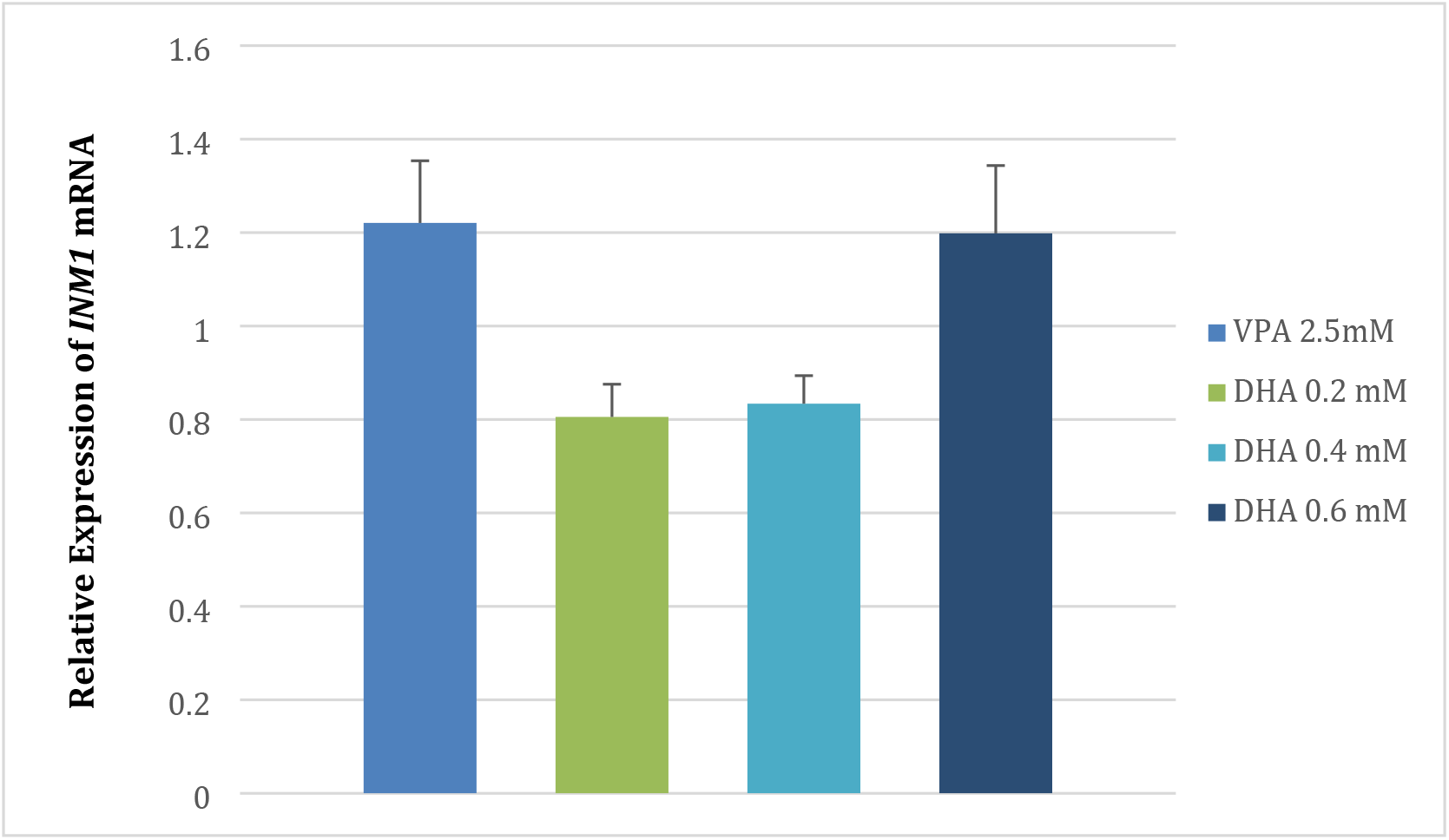
Relative *INM1* gene expression of cells grown in DHA (n =5) and positive control (VPA, n =5). The delta-delta CT method was used to determine fold change. *p* > 0.05 by ANOVA

## Discussion

Omega-3-fatty acids hold promise as natural treatments for bipolar disorder; however, more research is needed on their mechanism of action and effectiveness.

This study aimed to shed light on the molecular mechanism of action of omega-3-fatty acids, precisely the effect on the *myo*-inositol biosynthetic pathway, which has been implicated in the pathophysiology of bipolar disorder. This study represents the first to show the effects of any omega-3-fatty acid on intracellular *myo*-inositol and inositol biosynthesis genes. Employing *myo*-inositol perturbance as a screening tool, we showed that, unlike VPA, which has previously been shown to decrease intracellular *myo*-inositol, DHA resulted in more than a two-fold increase in *myo*-inositol compared to control (Figure 1). This suggests a different mode of action of DHA in alleviating bipolar disorder symptoms.

Both *INO1* and *INM1* belong to a class of genes containing *myo*-inositol-sensitive upstream activating sequence (UAS_ino_,) in their promoters. Not all UAS_ino_ containing genes have the same response to *myo*-inositol. For example, *INO1* is down-regulated by *myo*-inositol, while *INM1* is up-regulated by *myo*-inositol (Vaden et al., 2001; Murray and Greenberg, 2000). In this study, we showed relative expression of both genes is increased in the presence of DHA. Additionally, the increasing intracellular *myo*-inositol associated with growth in increasing DHA concentrations (Figure 1) was accompanied by declining *INO1* expression (Figure 2) and increasing *INM1* expression (Figure 3). The observed upregulation of *INM1* in the presence of VPA could be due to the presence of a second *myo*-inositol producing gene- *myo*-inositol monophosphate 1-phosphatase (*INM2*), also found in yeast.

We suggest a possible model for DHA’s effect on the inositol biosynthesis-DHA increases relative expression of *INO1* and *INM1*, resulting in increased production of *myo*-inositol. This may be accomplished via the *INO2* and *INO4* regulatory genes known to mediate the *myo*-inositol response of UAS_ino_ containing genes.

Lower levels of *myo*-inositol have been detected in postmortem brain samples of suicide victims and bipolar disorder patients compared to that of normal patients, while higher levels were observed in individuals in the manic phase (Shimon et al., 1997; Frey et al., 2005). Thus, drugs that deplete *myo*-inositol may be more effective in treating the manic phase, while the depressive phase may be more responsive to drugs that increase *myo*inositol. This is supported by reports that indicate lithium and valproate both of which lower intracellular *myo*-inositol levels, are more effective in treating the manic versus the depressive phase of bipolar disorder (Geddes et al., 2004; Geddes and Miklowitz, 2013). Meta-analyses suggest adjunctive use of omega-3 fatty acids improve the symptoms of depression but not those of mania (Sarris et al., 2012). We, therefore, suggest DHA exerts its therapeutic effect on bipolar disorder by increasing expression of *myo*-inositol biosynthetic genes resulting in increased intracellular *myo*-inositol during the depressive phase when levels are low and have little or no effect on mania when intracellular *myo*inositol levels are high. The implications of these findings are very significant since many of the currently approved anti-bipolar drugs have been shown to have harmful side effects, a problem less likely to occur with omega-3 fatty acids. Therefore omega-3 fatty acids may be an excellent alternative or at least an addition to current anti-bipolar medications and warrant further investigation.

